# On numerical study of Parkinson tremor

**DOI:** 10.1101/065318

**Authors:** Annalakshmi Harikrishna, Dennis Effah Osei, Magdalena Weronika Kamińska, Sarangam Majumdar

## Abstract

Parkinson’s disease and Parkinson’s tremor are the two most common movement disorders, nor do we fully understand the origin of one of the disease’s cardinal symptom: the Parkinsonian tremor. We study one mathematical model involved in Parkinson’s disease and in the Parkinsonian tremor. In this paper, we use the Van der Pol equation to further understand this tremor as well as investigate different numerical approaches to solve the system and compare them.

**AMS Subject Classification** 34L16; 34K28; 92B05

## 1 Introduction

Parkinson disease (PD) is a currently incurable progressive neurodegenerative disease identified by bradykinesia, stiffness, postural instability, which occurs in the later stages of PD, and rest tremors, often first seen in the hands or feet. Rest tremors are the most recognisable symptom in patients with PD and unlike physiological tremors, with a frequency range of 8-12Hz, rest tremors have a slower frequency in the range of 4-6Hz [1]. Although the full cause of PD is still unknown, it is linked to a deficiency of the neurotransmitter, dopamine, and elderly patients suffering from PD have been found to have reduced neuronal density in the substantia nigra [1]. Rest tremors are proposed to be caused by either central or peripheral feedback mechanisms or a combination of both [2]. For the central mechanisms, it is proposed that abnormal signals due to oscillations in the central nervous system are transmitted to muscles and result in the tremors seen in patients with PD [3, 4], and for the peripheral feedback mechanisms, it is thought that neurons involved in the muscle stretch reflex become unstable and no longer dampen muscle oscillations, resulting in tremors [5]. Mathematical modelling has proven to be a useful tool in further understanding PD tremors and what may potentially cause them. Previously, a model proposed by [6] using the Van der Pol equation, [6] discussed the possibilities of destroying downstream facilitatory paths to reduce the rest tremors. It was some time later that deep brain stimulation was proposed to be a possible treatment, where electrodes were implanted in patients brains to modulate any abnormal neural signals [7]. It has since proven to be effective against rest tremors in patients where medication was no longer sufficient. [6, 8] used a second order, nonlinear equation with one parameter that measured the ratio of emotional excitement to inhibition for their model. This model was proposed due to the observation in patients that tremors increased with excitation. Excitation has also been seen to affect bradykinesia, where patients are able to move faster at higher levels of excitation [1]. The Van der Pol equation was first proposed by Balthazar van der Pol (1889-1959), who was a Dutch electrical engineer who initiated modern experimental dynamics in the laboratory during the 1920’s and 1930’s. He first introduced his equation in order to describe triode oscillations in electrical circuits in 1927. The mathematical model has since become a well known second order ordinary differential equation with cubic nonlinearity, where thousands of articles have been published achieving better approximations to the solutions occurring in such non-linear systems. The Van der Pol oscillator is a classical example of a self-oscillatory system and is now considered to be a very useful mathematical model that can be used in much more complicated and modified systems. During the first half of the twentieth century, Balthazar van der Pol pioneered the fields of radio and telecommunications [9, 10, 11, 12, 13]. In an era when these areas were much less advanced than they are today, vacuum tubes were used to control the flow of electricity in the circuitry of transmitters and receivers. Contemporary with Lorenz, Thompson, and Appleton, in 1927, van der Pol experimented with oscillations in a vacuum tube triode circuit and concluded that all initial conditions converged to the same periodic orbit of finite amplitude. Since this behaviour was different from the behaviour of solutions of linear equations, van der Pol proposed a nonlinear differential equation commonly referred to as the unforced Van der Pol equation, as a model for the behaviour observed in the experiment. He also discovered the importance of the relaxation oscillations [11, 12]. The relaxation oscillations have become the cornerstone of the germetric singular perturbation theory and play a significant role in analysis. Moreover, it was noted that irregular noise was also appearing before transition from one subharmonical regime to another [13]. It was one of the first observations of chaotic oscillations in the electronic tube circuit. Since then it has been used by scientists to model a variety of physical and biological phenomena. In this paper, we extensively study this renowned equation numerically to obtain a better understanding of the Parkinsonian tremor. Moreover, we compare the different stiff and non-stiff ODE solvers according to their performance.

## 2 Mathematical Model

The Van der Pol oscillator is a model nonlinear differential equation where *p* is the position coordinate, which is a function of time, and *µ* ≥ 0 is a scalar parameter indicating the nonlinearity and the strength of the damping. The governing differential equation is

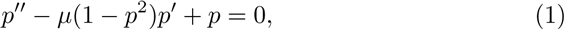

*µ* represents the emotional excitation and is defined as in [6] as the ratio of excitation (F) over inhibition (I)

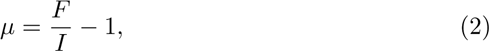

When *µ* = 0, meaning the level of excitation is equal to the level of inhibition, this is a simple harmonic oscillator and solutions have the form

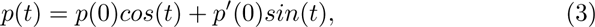

When *µ* is nonzero, the picture gets slightly more complicated. We can relate this as a class of ODE such as 

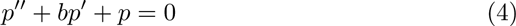

which have decaying oscillating solutions for *b* > 0 and exponentially growing solutions for *b* < 0. The coefficient *b* is interpreted as damping (with *b <* 0 corresponding to anti-damping behaviour where solutions gain energy over time). In the case of the Van der Pol equation, *b* is replaced by a nonlinear term which is negative when |*p*| < 1 and positive when |*p*| > 1. Therefore, the behaviour seen in practice is that there is a balance between growth behaviour for smaller *p* and decay behaviour for larger *p*. The result is that the solution bounces back and forth between slow motions for *p* > 1 and *p* < − 1 with fast transitions between, and the speed of those transitions is governed by the magnitude of *µ*.

Now equation (1) can be written as the dynamical system,*y*′ = *Ay*, with*y* (0) = *y*_0_. Solving this dynamical system, we can find that the eigenvalues of the Jacobian are*−* 1 and λ. We now analyze the system numerically and tally the results.

## 3 Numerical Experiment

In this section, we are numerically simulating the Van der Pol equation. We use Matlab to find the oscillatory behaviour. First, we took an example problem *y*′ = λ*y* with *y*(0) = 1 and λ = −10 and used the matlab code for the Euler Scheme to find the oscillatory behaviour with different step sizes. Figure 1 to Figure 4 shows the oscillations with the step size *h* = 0.184, 0.2, 0.3, 0.5. We observed that a small step size resulted in more oscillations than a large step size.

**Figure 1:**
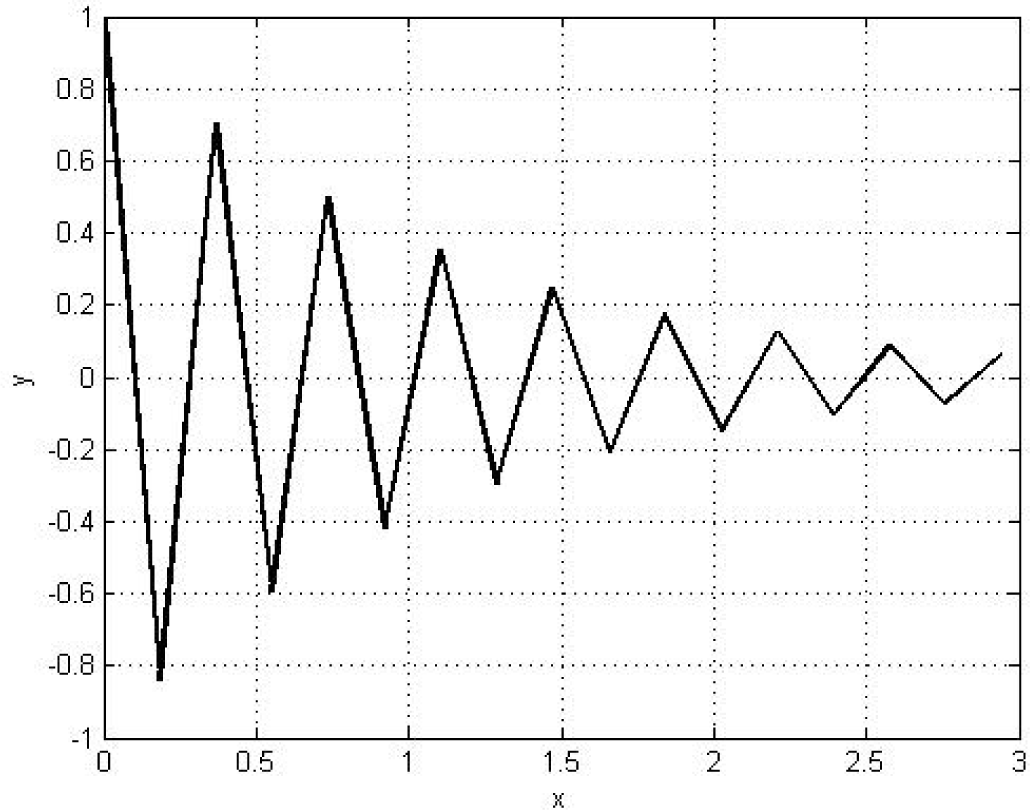
Oscillating behaviour for step size h = 0.184

**Figure 2:**
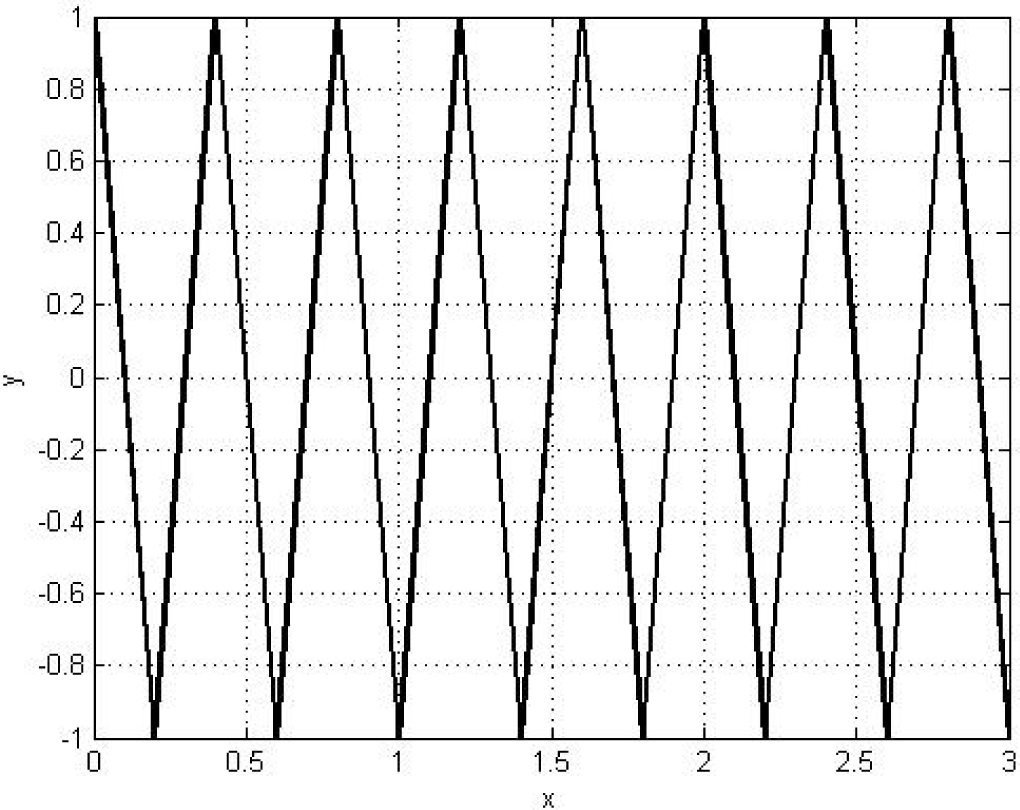
Oscillating behaviour for step size h = 0.2

**Figure 3:**
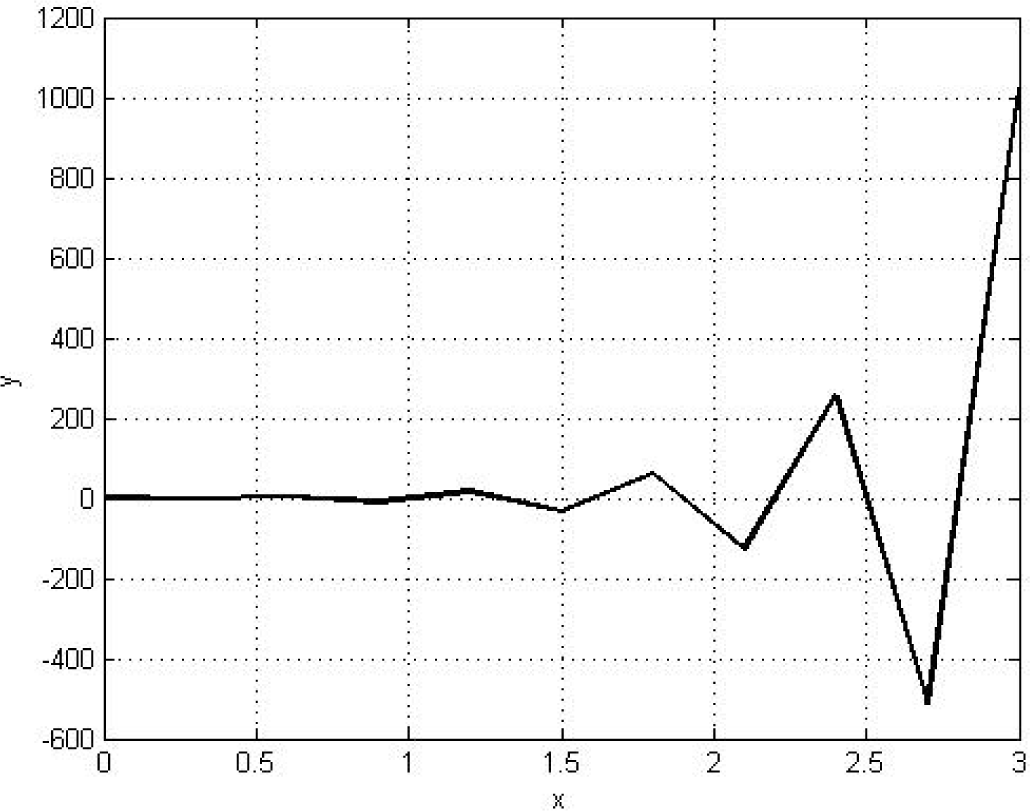
Oscillating behaviour for step size h = 0.3

**Figure 4:**
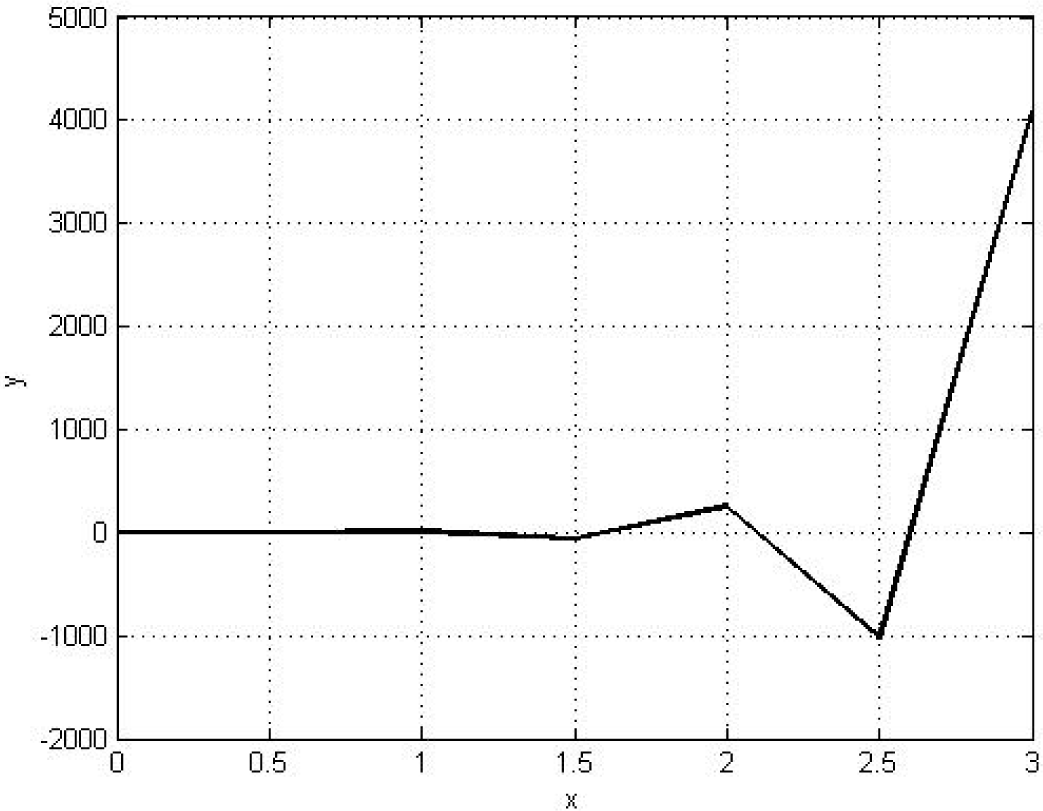
Oscillating behaviour for step size h = 0.5

### 3.1 Numerical Simulation of Van der Pol equation

We used the ODE45 and ODE15s solvers and compared the results. ODE45 is a non-stiff solver for the differential equations with a medium order method. It is based on the Runge-Kutta explicit scheme. On the other hand, ODE15s is a stiff solver with various order methods.

- For *µ* = 1 We found for ODE45.
  - 55 successful steps.
  - 8 failed attempts.
  - 379 function evaluations.
  - Elapsed time is 0.023931 seconds.
  For ODE15s
  - 193 successful steps.
  - 31 failed attempts.
  - 406 function evaluations.
  - Elapsed time is 0.087537 seconds.
  - 1 partial derivative.
  - 57 LU decompositions.
  - 402 solutions of linear systems.

Therefore, we can say that for *µ* = 1, ODE45 is the better choice due to it taking less time to compute and having a faster rate of convergence than ODE15s. We observe this because *µ* = 1 is a non-stiff equation. Figure 5 and Figure 6 illustrate solutions for both the solver and phase plane respectively.

**Figure 5:**
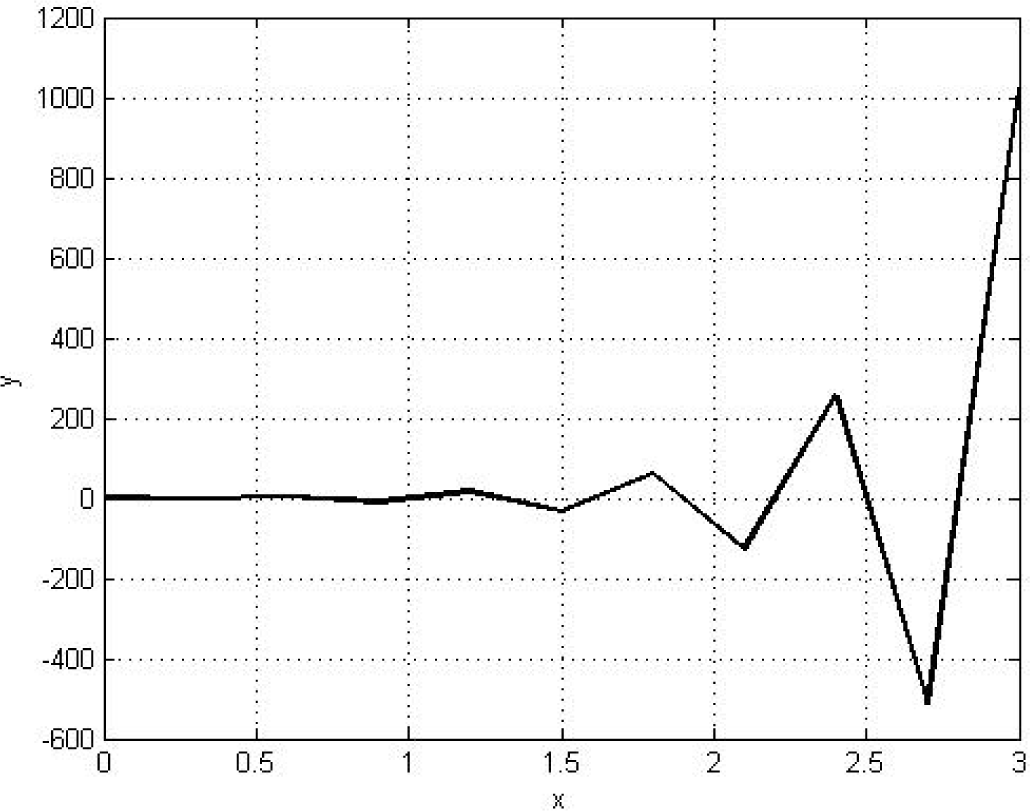
Oscillatory behaviour with time for *µ* = 1

**Figure 6:**
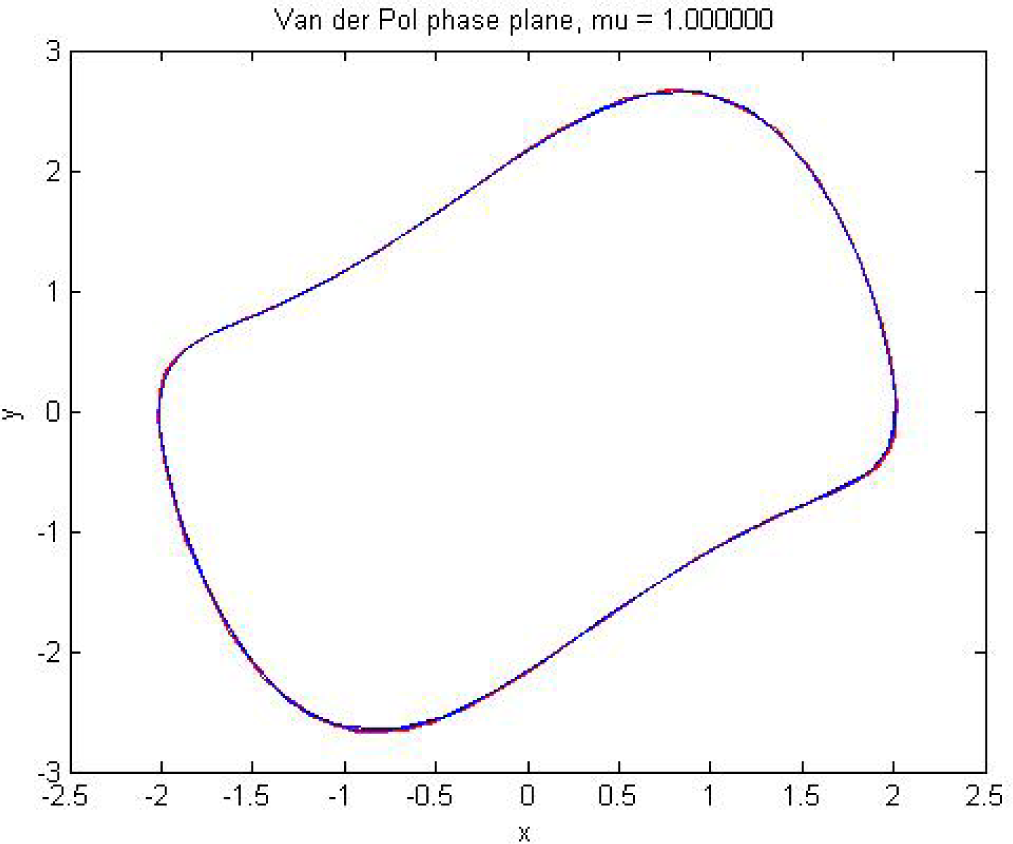
Phase plane for *µ* = 1

- For *µ* = 10 We found for ODE45.
  - 382 successful steps.
  - 55 failed attempts.
  - 2623 function evaluations.
  - Elapsed time is 0.044707 seconds.
  For ODE15s
  - 536 successful steps.
  - 138 failed attempts.
  - 1475 function evaluations.
  - Elapsed time is 0.159399 seconds.
  - 16 partial derivatives.
  - 192 LU decompositions.
  - 1426 solutions of linear systems.

Here, we see that ODE45 is also a better solver than ODE15s in all aspects for *µ* = 10. Figure 7 and Figure 8 show the solution curve and phase plane for *µ* = 10.

**Figure 7:**
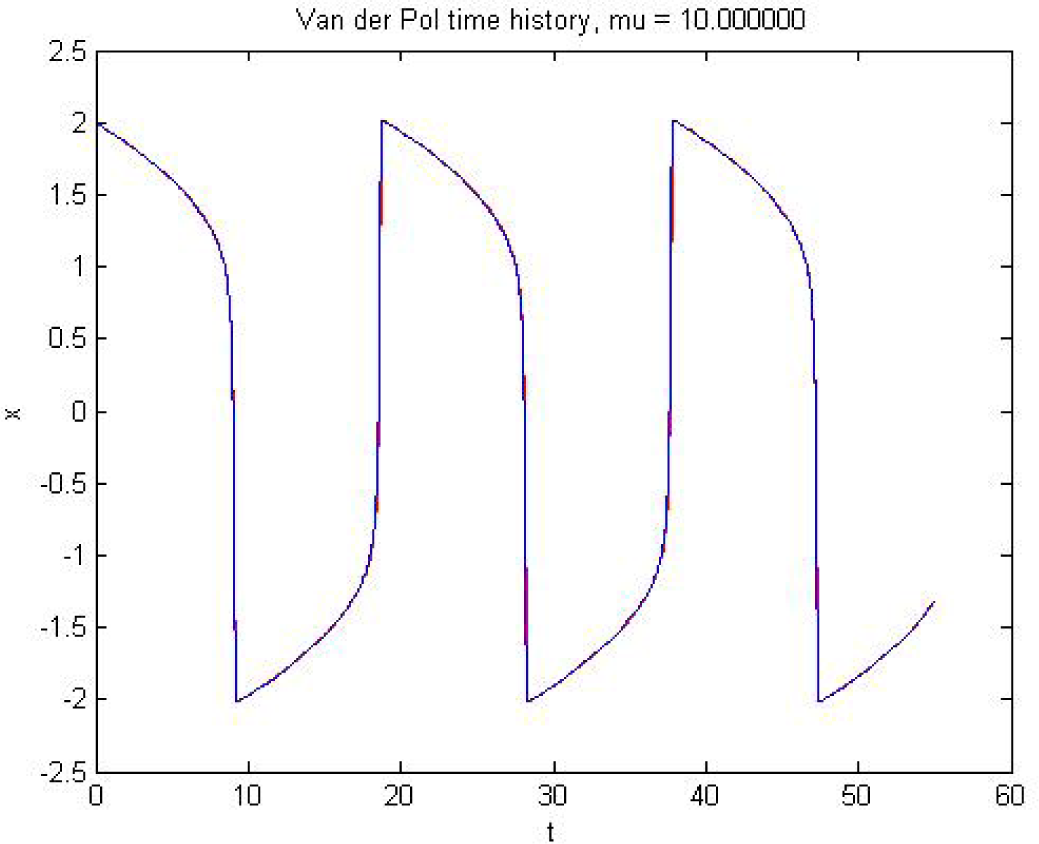
Oscillatory behaviour with time for *µ* = 10

**Figure 8:**
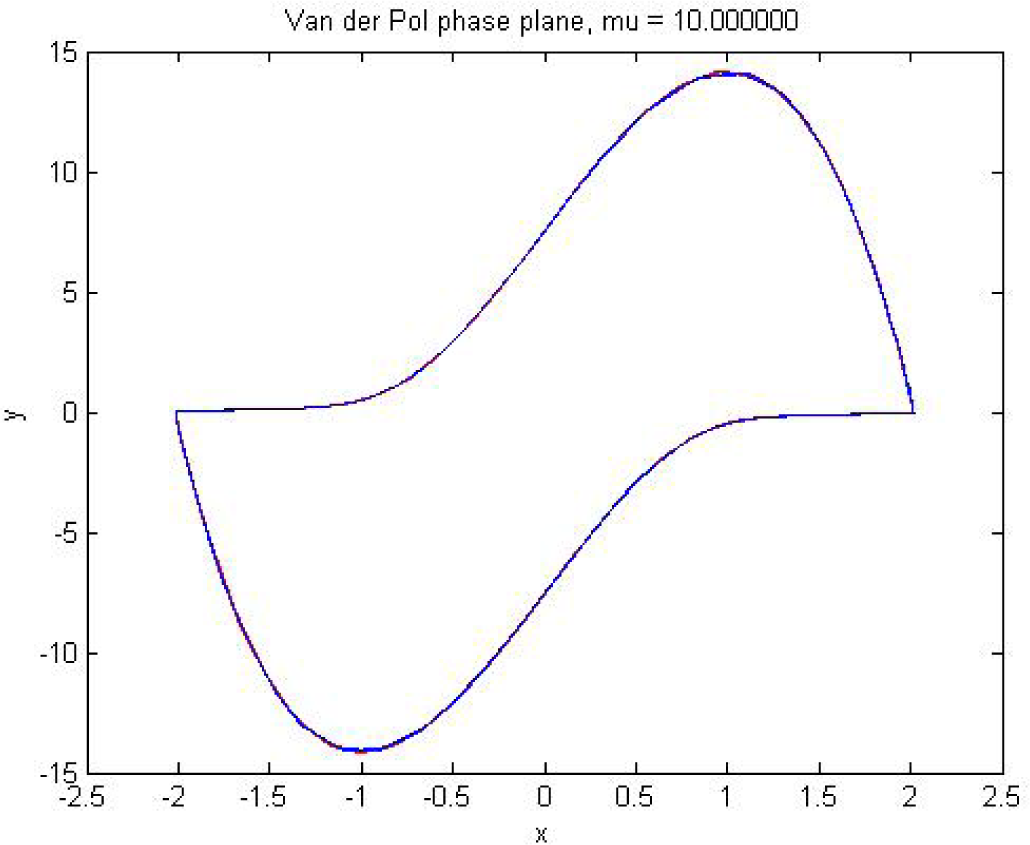
Phase plane for *µ* = 10

- For *µ* = 100 We found for ODE45.
  - 26368 successful steps.
  - 1701 failed attempts.
  - 168415 function evaluations.
  - Elapsed time is 2.574692 seconds.
  For ODE15s
  - 761 successful steps.
  - 282 failed attempts.
  - 2434 function evaluations.
  - Elapsed time is 0.251495 seconds.
  - 53 partial derivatives.
  - 357 LU decompositions.
  - 2274 solutions of linear systems.

Finally, we observe that ODE15s is a better solver than ODE45 because now the problem is stiff (for *µ* = 100). Both the solvers return results that are nearly indistinguishable visually. Figure 9 and Figure 10 portrait the nature of the solution and phase plane for the stiff problem.

**Figure 9:**
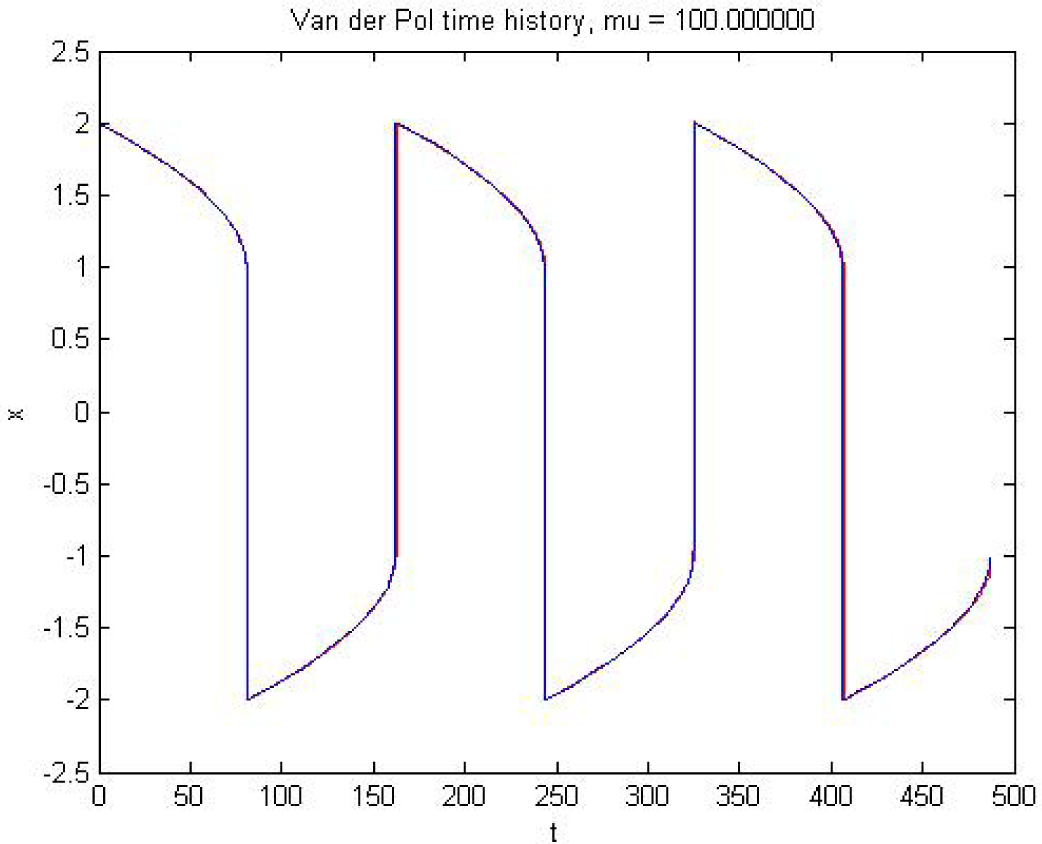
Oscillatory behaviour with time for *µ* = 100

**Figure 10:**
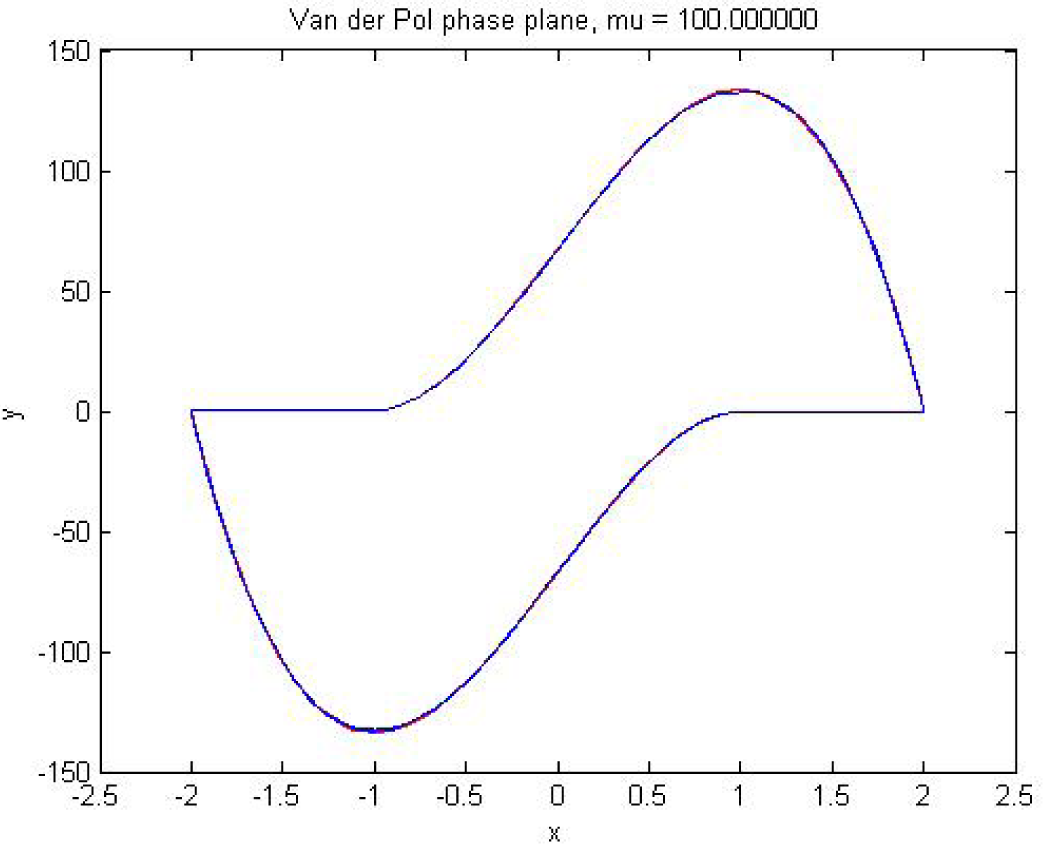
Phase plane for *µ* = 100

Figure 11 and Figure 12 show the oscillatory behaviour of the Van der Pol equation with two different step sizes. The smaller step size shows more oscillation.

**Figure 11:**
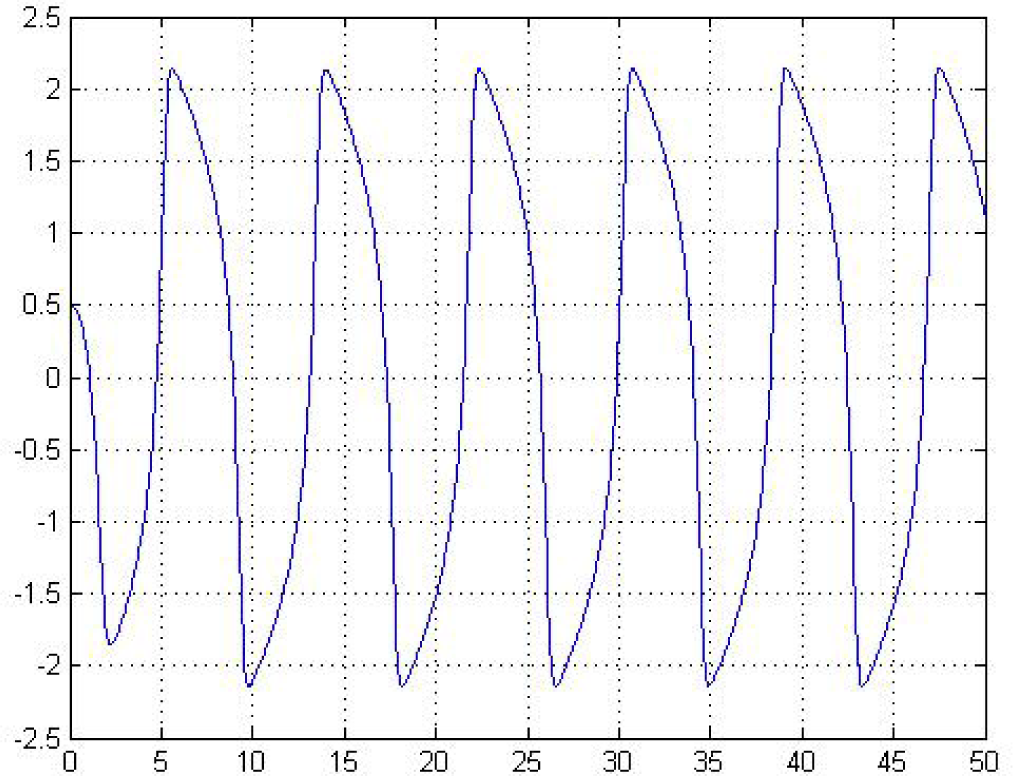
Oscillatory behaviour for step size h = 0.05

**Figure 12:**
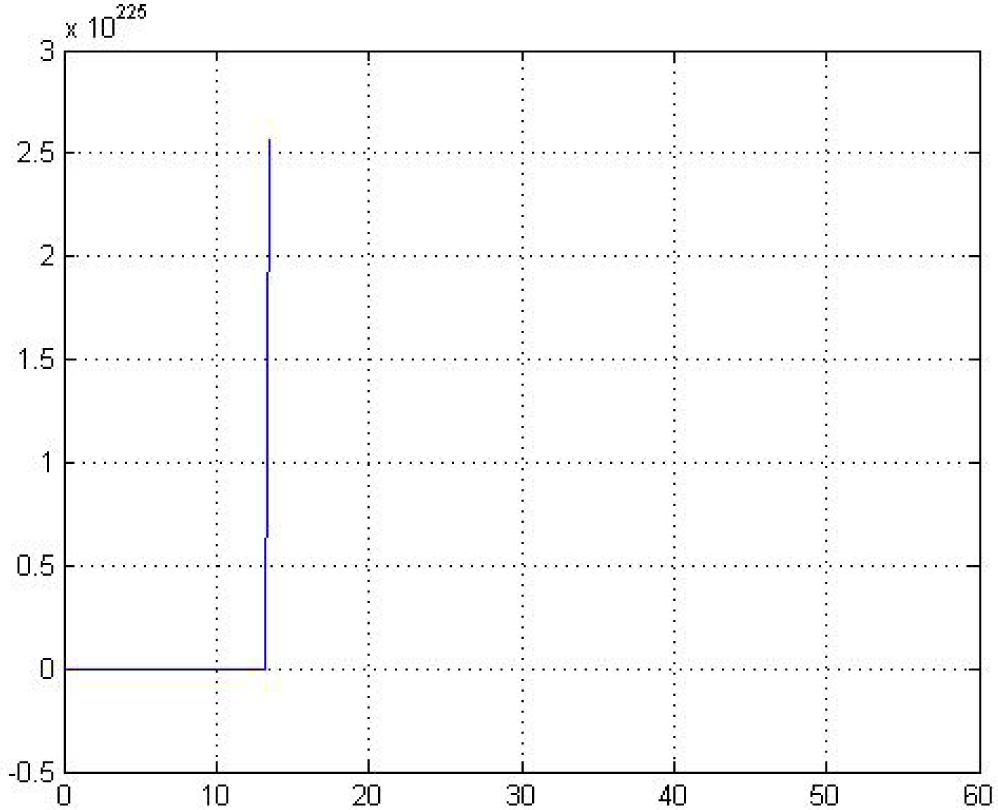
Oscillatory behaviour for step size h= 0.184

## 4 Conclusions

We studied the Van der Pol equation as a model of the Parkinson’s tremor and reported the numerical results. From these results, we can conclude that ODE45 works well for non-stiff problems while ODE15s performs better in the case of stiff problems. If we discretise the domain with a small step size, we obtain a good solution with less error. The model we have studied only considers the impact of emotional excitation rather than pinpointing the origin of the Parkinsonian tremor.

